# Equalizing Effect of Pollinator Adaptive Foraging on Plant Network Coexistence

**DOI:** 10.1101/2025.03.26.645069

**Authors:** Fernanda S. Valdovinos, Taranjot Kaur, Robert Marsland

## Abstract

The role of pollination in plant coexistence remains contested, with studies using Lotka-Volterra models suggesting pollination hinders while consumer-resource approaches indicating it can promote plant coexistence. Through the lens of modern coexistence theory, we analyze how network nestedness and pollinator foraging behavior influence plant coexistence via stabilizing (increasing the relative effect of intraspecific to interspecific competition) or equalizing (reducing fitness differences) mechanisms. Our findings reveal that while nestedness alone threatens specialist plant persistence, adaptive foraging by pollinators generates equalizing mechanisms by benefiting specialist over generalist plants which promotes plant coexistence, helping explain the contrasting conclusions from previous studies.

---

The role of mutualistic interactions—particularly pollination—in species coexistence remains a fundamental yet debated question in ecology^1–7^. Studies examining how pollination interactions influence plant coexistence have yielded contrasting results across different approaches. A theoretical analysis using empirically-parameterized Lotka-Volterra competition models in two-species systems suggest that pollination interactions generally hinder coexistence by favoring abundant species^7^. However, consumer-resource models incorporating pollinator behavior showed that adaptive foraging promotes plant coexistence in nested plant-pollinator networks through pollination niche partitioning^2–4^. While generalist plants typically outcompete specialists under fixed foraging patterns, adaptive foraging enables specialist plants to persist by receiving focused visitation from generalist pollinators, increasing the quantity and quality of visits specialists receive (Box 1). Supporting this plant coexistence mechanism, an empirical study of a highly modular plant-pollinator network found that rarer specialist plants actually benefit from more efficient pollination through reduced heterospecific pollen transfer^6^, revealing how pollination interactions can enhance plant coexistence by favoring specialists over generalists through pollinator niche partitioning.

These contrasting findings reflect fundamental differences in how each study conceptualizes plant-pollinator systems. Johnson et al^7^ parameterize Lotka-Volterra competition models using empirical measurements of seed production across varying neighbor densities, comparing scenarios from ambient-plus-supplemental-hand-pollination plants (reflecting only resource competition) versus ambient-pollinated plants (capturing both resource and pollinator-mediated competition)—an approach that sidesteps explicit representation of pollinators and their behaviors. In contrast, Valdovinos and collaborators^2–4^ demonstrate how pollinator adaptive foraging can generate niche partitioning through consumer-resource dynamics (Box 1), but their work did not explicitly connect these mechanisms to modern coexistence theory —as Johnson et al did— leaving unclear whether this niche partitioning generates stabilizing or equalizing (see definitions below) effects on plant coexistence. Finally, Wei et al^6^ used field observations and statistical modeling to link plant rarity, pollinator specialization, and pollination efficiency, showing that specialized interactions benefit rare plants through reduced heterospecific pollen transfer. However, like Valdovinos and collaborators, their work did not frame these findings within modern coexistence theory, leaving open how these specialized interactions translate into coexistence mechanisms that could maintain plant species diversity.

Here, we analyze Valdovinos et al.’s consumer-resource framework (Box 1) through the lens of Chesson’s modern coexistence theory to reconcile these contrasting findings from pairwise Lotka-Volterra models, mechanistic network consumer-resource models, and network field observations; as this framework has common components with all three approaches. We calculated plant coexistence as the fraction of initial plant species that coexisted (i.e., persisted) until the end of the simulations (see Methods) and used the mutual invasibility criterion by Chesson and Kuang^8^ to determine whether stabilizing or equalizing mechanisms are at play:

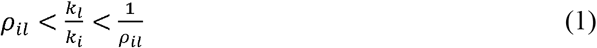

Where (*ρ*_*ik*_)is niche overlap between plant species *i* and *l*, and (*k*_*i*_) is *i*’s fitness. Stabilizing mechanisms increase negative intraspecific interactions relative to negative interspecific interactions via increasing niche differentiation^9^(1− *ρ*_*il*_), while equalizing mechanisms reduce relative fitness (*k*_*i*_/*k*_*l*_. Supplementary results (Eqs. S1-S13) show our derivations of *ρ*_*il*_ and *k*_*i*_ for Valdovinos et al’s model.

We found that nestedness decreases while adaptive foraging increases plant coexistence (Fig. 1A), operating through equalizing rather than stabilizing mechanisms. Neither factor affected niche overlap (*ρ*_*il*_), which remained unchanged across simulations with fixed and adaptive foragers (Fig. 1B) and between non-nested and nested networks (not shown). Instead, their influence manifested through plant fitness (Fig. 1C). With fixed foragers, generalist plants (visited by many pollinator species) showed markedly higher fitness than specialists (visited by one or a few pollinator species), particularly in nested networks. Adaptive foraging eliminated these fitness disparities, counteracting the amplifying effect of nestedness on generalist-specialist differences. Indeed, it nullified all positive correlations between plant fitness and plant generality measured as degree (number of pollinator species visiting the plant), as evidenced by near-zero slopes and R^2^ values in Table S1.

**Fig. 1.**
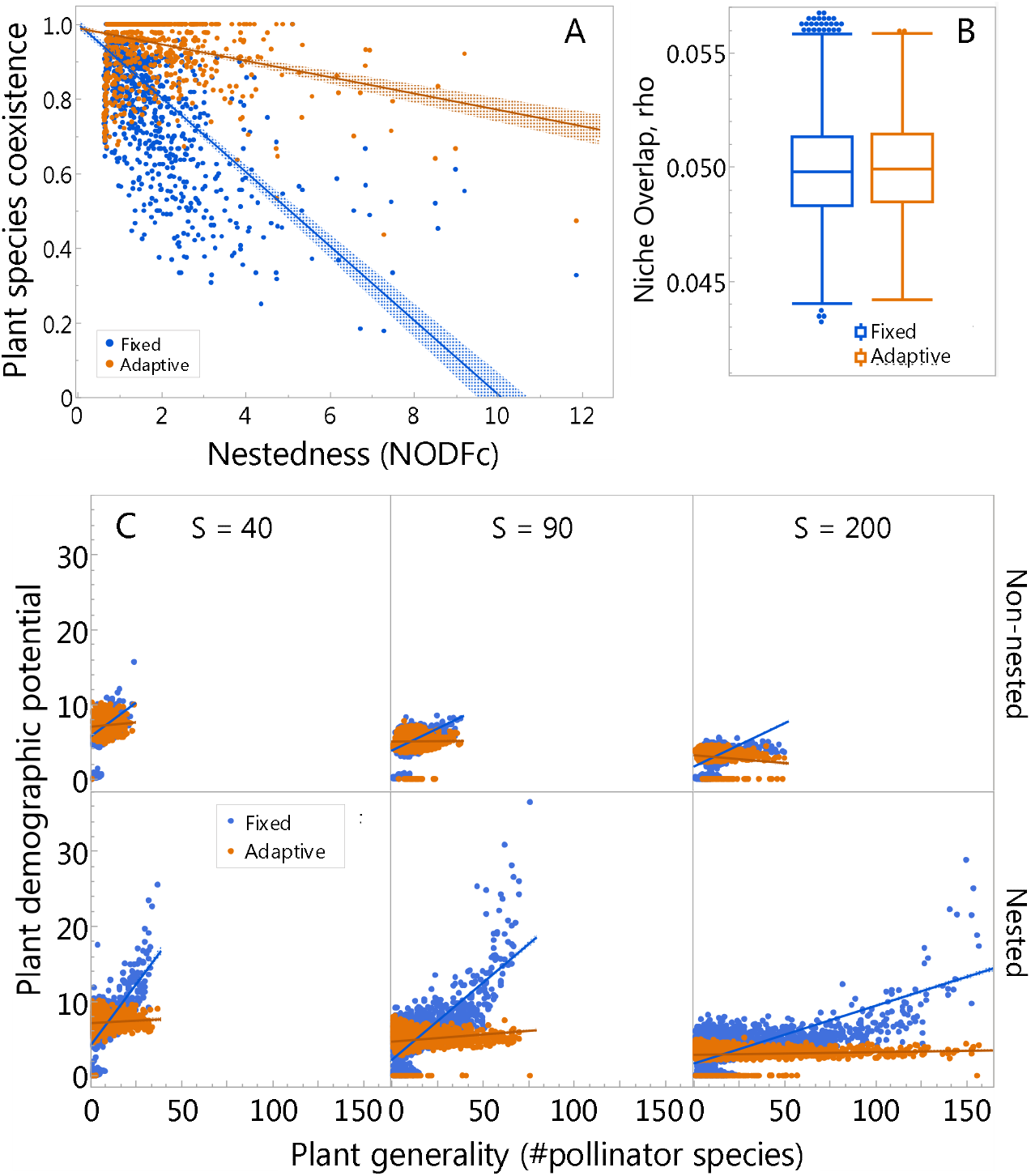
Nestedness decreases and adaptive foraging increases plant coexistence via equalizing mechanisms. **A)** Adaptive foragers weaken the strong negative relationship between plant coexistence and nestedness, measured as NODFc. Each dot represents plant coexistence (i.e., the fraction of plant species that persisted until the end of the simulations) for one of the 1200 networks with fixed (orange) and adaptive (blue), with 2400 total. **B)** Adaptive foragers do not affect niche overlap (*ρ*_*ik*_), as indicated by the similarity between the orange and blue box plots. Box plots display the median (middle line), the interquartile range (box), and the 95% confidence intervals (error bars) of the mean niche overlap across all plant species in all simulations. **C)** Nestedness exacerbates and adaptive foragers eliminate differences in demographic potential between generalist and specialist plants. Dots represent plant demographic potential (Eq. 2) for each plant species in each simulation against its degree (i.e., number of pollinator species visiting it), separated by network richness (S) and whether the network is non-nested or nested. Lines correspond to linear regressions used as reference to show tendencies. Regressions for fixed foragers have slopes between 0.1 and 0.6 and R^2^ between 0.15 and 0.6, while all regressions for adaptive foragers have near zero slopes and R^2^. See Table A1 for all statistics.

To better understand how this equalizing mechanism operates, we examined the fitness expression (derived in Eqs. S10-S13) representing the difference between per-capita seed recruitment rate to adults (*g*_*i*_*e*_*i*_*v*_*i*_) and plant mortality rate 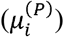:

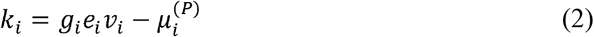

where *g*_*i*_, *e*_*i*_, and 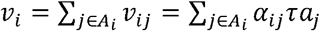 represent *i*’s maximum recruitment rate, seed production, and total per-plant visits received, respectively. The per-plant visits from pollinator *j* (*v*_*ij*_) depends on *j*’s foraging effort (*α*_*ij*_), efficiency (*τ*), and density (*α*_*j*_). This equation reveals a direct linear relationship between demographic potential and total per-plant visits, demonstrating that nestedness and adaptive foraging affect demographic potential through their effects on total per-plant visits. Indeed, mirroring Fig. 1C, nestedness amplifies visit disparities between generalist and specialist plants (Fig. S1) while adaptive foraging equalizes them (Fig. 1C), eliminating the positive correlation between plant degree and total per-plant visits (Table S2).

This coexistence mechanism of equalizing total per-plant visits becomes even clearer when examining the expression of plant density at equilibrium (derived in Eq. S14-S19):

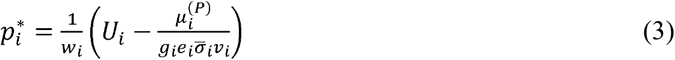

where,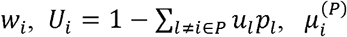, and 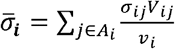are *i*’s intra-specific competition coefficient for recruitment, recruitment potential discounting for total inter-specific competition, mortality rate, and weighted average visit quality, respectively. Note that 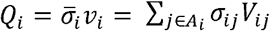 represents *i*’s total pollination events (see Methods). Eq. 3 indicates that sufficiently high pollination events decrease the negative effect of plant mortality on abundance to zero, allowing plants to reach their maximum abundance, which is determined by seed recruitment. More importantly for plant coexistence, Eq. 3 leads to a threshold of pollination events required for plant persistence:

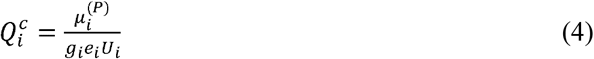

When 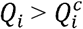, plant species *i* persists and reaches its equilibrium (Eq. 3). When 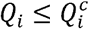, plant species *i* goes extinct, reaching the other equilibrium (i.e.,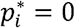). That is, plant species that do not receive that critical amount of pollination events, go extinct, which happens either because all their pollinator species went extinct or because their pollinators did not assign enough foraging effort to them (as a result of the plants offering less rewards than the other plants in the pollinators’ diet).

Our findings show that modern coexistence theory classifies niche partitioning driven by adaptive foraging^2–4^ (Box 1) as equalizing rather than stabilizing. Two reasons explain this discrepancy with the typical assumption of niche partitioning as a stabilizing mechanism^9^. First, adaptive foraging-driven niche partitioning increases specialist plant fitness to match generalists’, making it an equalizing effect. Second, this consumer-resource model assumes stronger intraspecific than interspecific effects in seed recruitment competition—a strong stabilizing mechanism operating at a longer timescale than pollination. This long-term stabilizing mechanism in seed recruitment counteracts pollination’s short-term effects on Chesson’s niche overlap (*ρ*_*ik*_), rendering pollination’s influence negligible. Fig. 2D,C (in Box 1) demonstrates seed recruitment’s counteracting effect on pollination, showing decreased seed recruitment when species produce abundant seeds due to increased pollination.

**Fig. 2.**
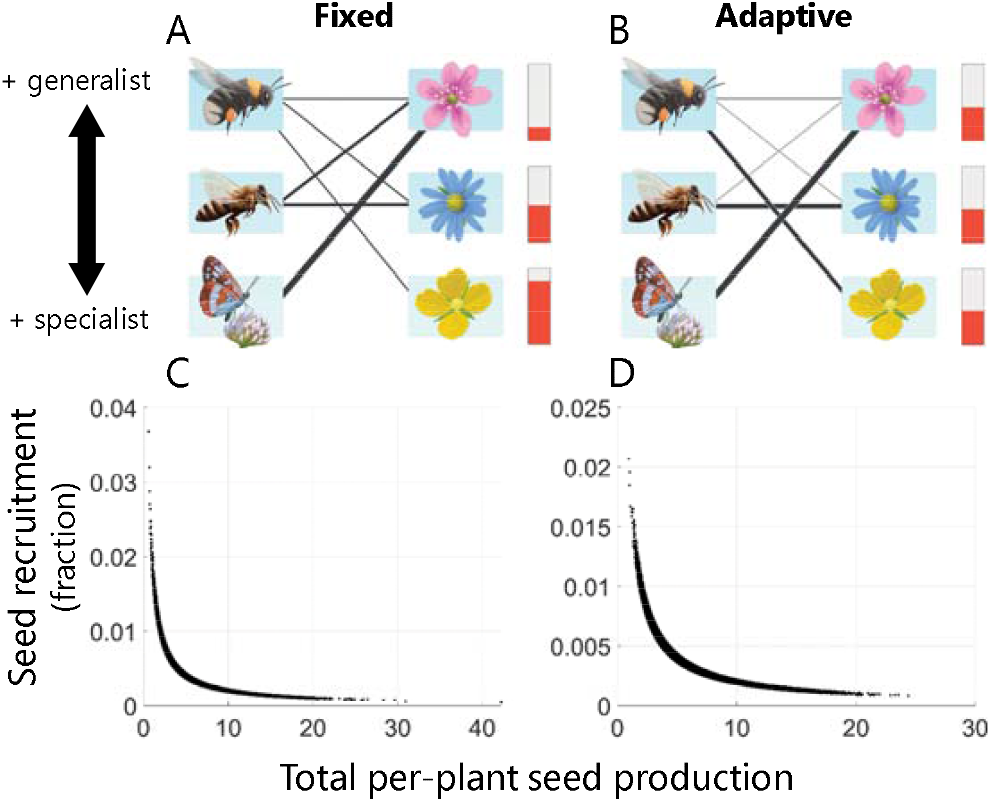
(A) Fixed foraging scenario where pollinators distribute foraging efforts equally among their plant partners (illustrated by line thickness), which results in generalist and specialist plants receiving many and very few visits, respectively. This visit distribution, in turn, results in rewards of generalist plants being very depleted while those of specialist plants being abundant (red bars, Fig. S2). (B) Adaptive foraging scenario where pollinators adjust their efforts based on available rewards, resulting in more focused visitation to specialist plants and equal final rewards among plant species (Fig. S2). Line thickness indicating visit frequency. (C, D) Inverse relationship between seed recruitment (as fraction of seeds) and total per-plant seed production shows how competition for seed recruitment strongly counteracts pollination effects on seed production both in the case of fixed (C) and adaptive (D) foragers.

Our analysis explains previous contrasting results. Johnson et al.^7^ compared plant competition coefficients under ambient versus ambient-plus-supplemental-hand-pollination, finding lower niche differences under ambient pollination. The lower niche differences predicted reduced plant coexistence under ambient pollination, suggesting animal pollination decreases plant coexistence. However, their model combined in one coefficient competition for both seed recruitment and pollinators. Our results suggest that combining these two processes can confound pollination effects on coexistence given the counteracting effect of seed recruitment on the effect of animal pollination on seed production (Fig. 2D,C). These competition sources operate in nature at different temporal scales—pollination competition occurs over hours or days, while seed recruitment competition spans seasons or years—and our model reflects this temporal separation. Furthermore, since their experimental design did not include measurements of pollinator visits or preferences, it cannot directly validate their proposed mechanism that pollinators decrease plant coexistence by preferring abundant plants. Instead, our results align with Wei et al.’s findings of strong pollinator niche partitioning, where rarer species showed greater pollinator specialization leading to higher pollination-mediated fitness than abundant species^6^. Wei et al.’s comprehensive network analysis demonstrated that specialist plants benefited from improved delivery of conspecific pollen and lower interference from heterospecific pollen transfer. This evidence for niche partitioning favoring rare specialist plants directly contradicts Johnson et al.’s untested mechanism of pollinators preferring abundant plants, and supports our findings of adaptive foraging enhancing plant coexistence via equalizing mechanisms.

## Box 1

**Consumer-Resource Framework for Plant-Pollinator Networks**

Valdovinos et al. model plant-pollinator interactions by decomposing the mutualistic relationship into its mechanistic components: pollinators consume floral rewards while serving as pollen vectors between conspecific plants during their foraging visits. Plant-pollinator networks typically display a nested structure, where specialist species interact primarily with generalist partners, while generalist species interact with both specialist and generalist partners. Within this consumer-resource framework, a nested structure leads to strong niche overlap in both guilds: plant species heavily overlap in their use of pollination services (sharing generalist pollinators), while pollinators overlap in their use of floral rewards from generalist plants (Fig. 2A-B).

Under fixed foraging (Fig. 2A), pollinators distribute their visits equally among their plant partners (*α*_*ij*_ =1/#*P*_*j*_, where #*P*_*j*_ is the number of plant species pollinator *j* visits). This strategy disadvantages specialist plants, as they receive fewer and lower-quality visits: only generalist pollinators visit them, and these pollinators allocate just a small fraction of their total visits to specialist plants, ultimately leading to their extinction. Under adaptive foraging, pollinators adjust their foraging efforts (*α*_*ij*_) based on the relative availability of floral rewards among plants in their diet. This leads generalist pollinators to concentrate their efforts on specialist plants that maintain higher reward levels due to receiving fewer visitors overall. This allows specialist plants to persist given that they receive more and higher quality visits.

## Methods

We use a consumer-resource model for simulating the dynamics of plant-pollinator networks ^2–4^, where nodes represent species and links indicate which pollinator species visits which plant species. This model defines the population dynamics of each plant *i* (*p*_*i*_, Eq. 1) and animal *j* (*a*_*j*_, i.e., pollinator, Eq. 2) species in the network, the dynamics of floral rewards made available to pollinators by each plant species *i* (*R*_*i*_, Eq. 3), and the foraging effort that each pollinator species *j* per-capita assigns to each plant species *i* in its diet (*α*_*ij*_, Eq. 4), as follows:

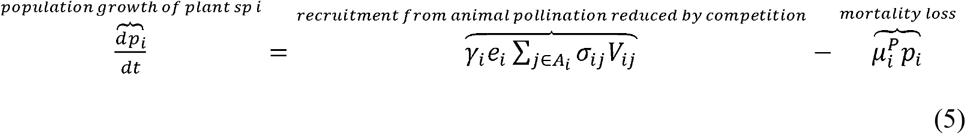

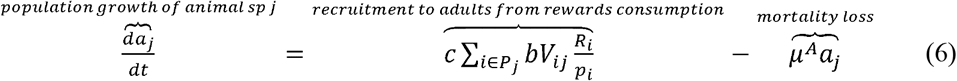

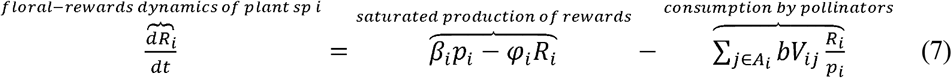

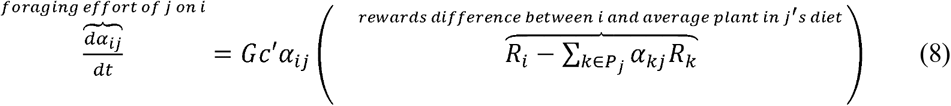

A key variable connecting these four equations is the visitation rate of pollinator species *j* to plant species *i*, defined as:

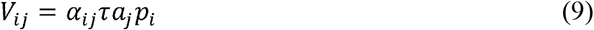

Where *τ* represents the flying efficiency of pollinator *j*. For simplicity, we assume all pollinators have the same efficiency *τ* =1, thus we dropped the subindex *j*. Plant *i*’s population grows with increased seed recruitment to adults and decreases with mortality loss which depends on the per-capita plant mortality rate, 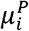. The recruitment term of Eq. 1, defines that only a fraction 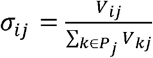 (i.e., visit quality of *j* on *i*) of the visits received from pollinator species *j* (*V*_*ij*_) become pollination events, then only a fraction of the total pollination events 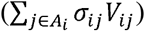 become seeds (where *A*_*i*_ indicates the set of pollinator species that visit plant *i*), and finally only a fraction *γ*_*i*_ of those total seeds recruit to adults, defined as:

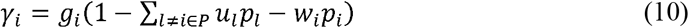

where *g*_*i*_ is the maximum fraction of total seeds that recruit to plants, and *u*_*l*_ and *w*_*i*_ are the inter-specific and intra-specific competition coefficients determining the competition strengths among plants for resources other than pollinators (e.g., soil, water, light). Inter-specific competition occurs among the set of all plant species in the network, *P*.

Pollinator population *j* grows (Eq. 2) with increasing recruitment to adults from rewards consumption and decreases with increasing mortality loss, where *μ*^*A*^ is the per-capita mortality rate for all animal species. Pollinator recruitment is determined by the rewards extracted in its visits to 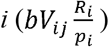 where *b* is the pollinator extraction efficiency of rewards in each visit which are converted to adults at a conversion rate *c*. We dropped the subindex *j* from all pollinator-specific parameters as we assumed — for simplicity given that our question is on plant coexistence not on pollinator variability — same parameter values across all pollinator species. Pollinators, however, vary drastically in their number of interactions given the network structure and in their resulting abundances. Floral rewards increase in a saturated manner following 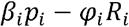, where *β*_*i*_ is the per-capita production rate of floral rewards and *φ*_*i*_ is self-limitation of rewards production, and decrease via pollinator consumption (Eq. 3). Finally, the foraging effort that *j* assigns on *i* increases when the difference 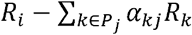 is positive, that is, when rewards of *i* are higher than rewards of an average plant in *j*’s diet 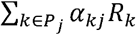 (Eq. 4) — where *P*_*j*_ is the set of plant species visited by *j* — and decreases when such difference is negative. The change in *α*_*ij*_ is faster with higher pollinator response rate, and with higher pollinator conversion and efficiency rates *c*’=*cbτ*. Foraging efforts are proportions between 0 and 1, and they sum to 1 for each pollinator species (i.e.,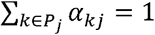). Thus, 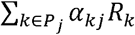 is mathematically equivalent to the definition of a weighted arithmetic mean. That is, a weighted average of the rewards, where the weights are the foraging efforts.

We use this model to simulate the dynamics of 1,200 algorithm-generated networks (Fig. 1A) without and with adaptive foraging, with a total of 2,400 simulations. Simulations without adaptive foraging assume fixed foraging efforts defined as *α*_*ij*_ = 1/*d*_*j*_, where *d*_*j*_ is the number of plant species that pollinator species *j* visits (i.e., pollinator *j*’s degree). Simulations with adaptive foraging assume that pollinators prefer plants that have more floral rewards available to them (i.e., their *α*_*ij*_ vary following Eq. 4).

The generated networks (Fig 1A) expand ranges of connectance (*C*) between 0.34 and 0.041 and species richness (*S*) between 15 and 238, with *C* × *S* combinations that resemble those observed in empirical networks (Fig. S1) and with an average ratio of pollinator to plant number of species of 2.5. Connectance is calculated as *C = L/(A×P)*, where *L* is the realized number of links, and *A×P* the potential links between the total number of pollinator species, *A*, and plant species, *P*, in the network, while *S* = *A+P*. These networks cover values from anti-nested to highly nested slightly beyond the ones observed in empirical networks to ensure encompassing the entire empirical range. We generated the networks using the algorithm developed by Thébault and Fontaine (2010). We calculated nestedness and its statistical significance using NODFst ^12^, using the R package bipartite ^13^. However, to compare across networks, we calculated NODFc ^14^ using R package maxnodf ^15^, which is known to better compare across networks of different sizes. We ran model simulations for 10,000 timesteps using Matlab R2024a (The MathWorks Inc.)

The mutual invasibility criterion of Eq. 5 was developed for pairwise interactions and indicates whether or not each species can invade a system where the other is present when starting at very low abundances ^8^. This criterion, however, accurately represented the coexistence between every 1,348,134 plant species pairs of our 2,400 simulations, except for 84 pairs involving the same species within a network in 6 networks without adaptive foraging. We tested that this was the case by evaluating the criterion at steady state (i.e., at the end of the simulations after 10,000 time-steps) for each plant species pair in each of our simulations and contrasting the outcome of Eq. 5 with whether the pair of species coexisted until the end of the simulations.

Note that we are using these calculations as metrics for describing coexistence mechanisms rather than as proof for coexistence given that our simulations can tell us for certain whether plant species coexisted or not.

## Supporting information

All supplemental tables and figures

All mathematical analyses

## Acknowledgments

This research was funded by National Science Foundation grant DEB-2129757.

## Data and Code Accessibility Statement

The code used to produce all results of this work is publicly available at GitHub repository https://github.com/Valdovinos-Lab/Plant_Coexistence.

## Author Contribution Statement

F.S.V. conceptualized, designed, and acquired funding for this study, run simulations, conducted analyses, created figures, and wrote the manuscript. T.K. conducted mathematical analyses in close collaboration with F.S.V and R.M.III. All authors reviewed the manuscript.

