## Supplementary material for "Equalizing Effect of Pollinator Adaptive Foraging on Plant Network Coexistence": All supplemental tables and figures

**List of Extended Data tables:**

1. Table S1. Regression statistics for demographic potential vs plant degree for all 1200 algorithm-generated networks separated by richness and nestedness levels, with fixed and adaptive foragers.
2. Table S2. Regression statistics for total visits per-plant vs plant degree for all 1200 algorithm-generated networks separated by richness and nestedness levels, with fixed and adaptive foragers.
3. Table S3. Model state variables, functions, and parameters.

**List of Extended Data figures:**

1. **Fig. S1. Adaptive foraging equalizes the visits that specialist and generalist species receive per plant.**
2. **Fig. S2. Final floral rewards with respect of plant species’ degree.**
3. **Fig. S3.** Connectance (C) vs species richness (S) of 155 empirical plant-pollinator networks obtained from <https://www.web-of-life.es/> that contain between 15 and 240 species**.**
4. **Fig. S4.** Connectance (C) vs species richness (S) of the 1200 algorithm-generated networks used in this study.

**Table S1.** Regression statistics for demographic potential vs degree for all 1200 algorithm-generated networks separated by richness and nestedness levels, with fixed and adaptive foragers. Slope and R^2^ are in bold to highlight that adaptive foragers eliminated the positive relationship between degree and demographic potential. Numbers in parentheses correspond to the standard error and t-statistic, respectively, of the equation estimate of the regression (slope or intercept).

| **Regression** | **S=40** | **S=90** | **S=200** |  |
| --- | --- | --- | --- | --- |
| **Demographic potential vs degree** with fixed foragers | **Slope: 0.18**  (0.008 \| 21.1)  Intercept: 5.6  (0.07 \| 2.3)  **R^2^: 0.15** | **Slope: 0.12**  (0.004 \| 27.5)  Intercept: 7.0  (0.05 \| 80.3)  **R^2^: 0.15** | **Slope: 0.11**  (0.003 \| 36.0)  Intercept: 1.7  (0.04 \| 46.3)  **R^2^: 0.15** | **Non-nested** |
| … with adaptive foragers | **Slope: 0.01**  (0.007 \| 2.29)  Intercept: 7.0  (0.054 \| 127.9)  **R^2^: 0.00**  N= 2609 | **Slope: -0.00**  (0.004 \| -0.06)  Intercept: 5.0  (0.04 \| 120.0)  **R^2^: 0.00**  N= 4302 | **Slope: -0.02**  (0.003 \| -4.8)  Intercept: 3.1  (0.04 \| 81.2)  **R^2^: 0.00**  N= 7440 |  |
| … with fixed foragers | **Slope: 0.32**  (0.01 \| 32.0)  Intercept: 4.0  (0.1 \| 42.1)  **R^2^: 0.39** | **Slope: 0.20**  (0.003 \| 63.6)  Intercept: 2.1  (0.04 \| 47.4)  **R^2^: 0.44** | **Slope: 0.08**  (0.001 \| 66.5)  Intercept: 1.6  (0.02 \| 76.1)  **R^2^: 0.26** | **Nested** |
| … with adaptive foragers | **Slope: 0.01**  (0.00 \| 2.5)  Intercept: 7.0  (0.05 \| 143.5)  **R^2^: 0.00**  N= 1588 | **Slope: 0.02**  (0.00 \| 10.6)  Intercept: 4.5  (0.03 \| 153.1)  **R^2^: 0.02**  N= 5171 | **Slope: 0.00**  (0.00 \| 5.1)  Intercept: 2.8  (0.01 \| 225.8)  **R^2^: 0.00**  N= 12897 |  |

**Table S2.** Regression statistics for total visits per-plant vs degree for all 1200 algorithm-generated networks separated by richness and nestedness levels, without and with adaptive foraging (AF).

| **Regression** | **S=40** | **S=90** | **S=200** |  |
| --- | --- | --- | --- | --- |
| **Total visits per-plant vs degree**  with fixed foragers | Slope: 0.56  Intercept: 17.6  R^2^: 0.18 | Slope: 0.36  Intercept: 11.78  R^2^: 0.17 | Slope: 0.36  Intercept: 5.1  R^2^: 0.16 | **Non-nested** |
| … with adaptive foragers | Slope: 0.04  Intercept: 21.9  R^2^: 0.00 | Slope: -0.00  Intercept: 15.8  R^2^: 0.00 | Slope: -0.05  Intercept: 9.7  R^2^: 0.00 |  |
| … with fixed foragers | Slope: 1.0  Intercept: 12.6  R^2^: 0.41 | Slope: 0.63  Intercept: 6.6  R^2^: 0.45 | Slope: 0.24  Intercept: 5  R^2^: 0.26 | **Nested** |
| … with adaptive foragers | Slope: 0.05  Intercept: 12.6  R^2^: 0.01 | Slope: 0.07  Intercept: 14  R^2^: 0.02 | Slope: 0.01  Intercept: 8.7  R^2^: 0.00 |  |

**Table S3. Model state variables, functions, and parameters.** Values were drawn from a uniform random distribution with the specified mean, and variances of 10% and 0% of means for plant and animal parameters, respectively. That is, all pollinator species had the same parameter values and, therefore, the subscript *j* was dropped for clarity. Parameter were taken from Valdovinos et al. (2013) and (2018). Asterisks indicate initial conditions. $P_{j}$ and $A_{i}$ are the subsets of plant species that *j* visits and animal species the visit plant *i*, respectively. $d_{j}$ is the degree of pollinator *j*, that is, the number of species *j* visits.

| Definition | Symbol | Dimension | Mean value |
| --- | --- | --- | --- |
| ***State Variables*** |  |  |  |
| Density of plant population *i* | $p_{i}$ | individuals area^-1^ | 0.5* |
| Density of animal population *j* | $a_{j}$ | individuals area^-1^ | 0.5* |
| Total density of floral rewards of plant population *i* | $R_{i}$ | mass area^-1^ | 0.5* |
| Foraging effort of *j* on *i* | $\alpha_{ij}$ | None | $\frac{1}{d_{j}}$ * |
| ***Functions*** |  |  |  |
| Visitation rate of *j* to *i* (quantity of visits) | $V_{ij}=\alpha_{ij}\tau a_{j}p_{i}$ | visits area^-1^ time^-1^ | variable |
| Quality of visits of *j* (per-capita) to *i* (per-capita) | $\sigma_{ij}=\frac{V_{ij}}{\sum_{k\in P_{j}} V_{kj}}$ | None | variable |
| Fraction of seeds *i* that recruit to adults | $\gamma_{i}=g_{i}\left( 1-\sum_{l\neq i\in P_{j}} u_{l}p_{l}-w_{i}p_{i} \right)$ | None | variable |
| ***Parameters*** |  |  |  |
| Visitation (flying) efficiency | $\tau$ | visits area time^-1^ individuals^-1^ individuals^-1^ | 1 |
| Expected number of seeds produced by a pollination event | $e_{i}$ | individuals visits^-1^ | 0.8 |
| Per-capita mortality rate of plants | $\mu_{i}^{(P)}$ | time^-1^ | 0.02 |
| Conversion efficiency of floral rewards to pollinator births | $c$ | individuals mass^-1^ | 0.2 |
| Per-capita mortality rate of pollinators | $\mu^{(A)}$ | time^-1^ | 0.001 |
| Pollinator extraction efficiency of rewards in each visit | $b$ | individuals visits^-1^ | 0.4 |
| Maximum fraction of total seeds that recruit to plants | $g_{i}$ | None | 0.4 |
| Inter-specific competition coefficient of plants | $u_{l}$ | area individuals^-1^ | 0.06 |
| Intra-specific competition coefficient of plants | $w_{i}$ | area individuals^-1^ | 1.2 |
| Production rate of floral rewards | $\beta_{i}$ | mass individuals^-1^ time^-1^ | 0.2 |
| Self-limitation parameter of rewards production | $\varphi_{i}$ | time^-1^ | 0.04 |
| Adaptation rate of foraging efforts of pollinators | $G$ | None | 2 |

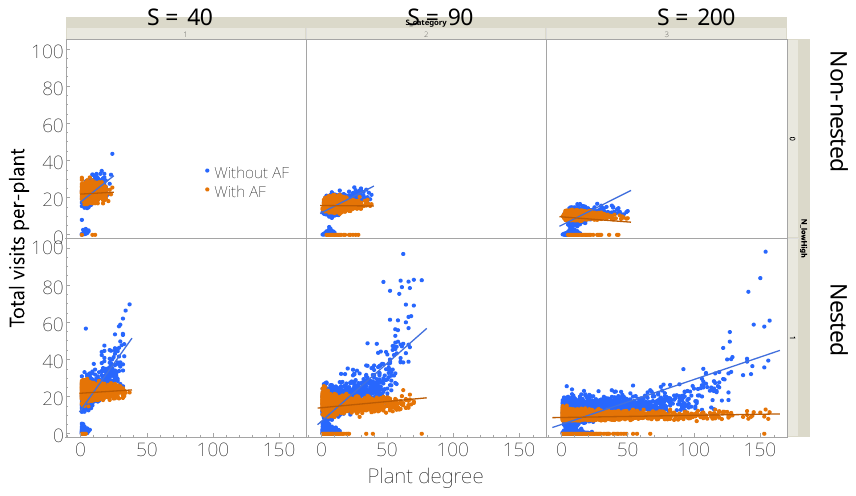

**Fig. S1.** **Adaptive foraging equalizes the visits that specialist and generalist species receive per-plant.** Dots represent total visits per-plant for each plant species in each simulation against its degree (i.e., number of pollinator species visiting it), for all simulations (i.e., three network richness levels and for nested and non-nested). Blue and orange dots correspond to simulations without and with adaptive foraging, respectively. All the statistics of the 12 regressions can be found in Table A2. Without AF, the slopes of the linear regressions range from 0.36 to 0.56 with R^2^ from 0.16-0.18 in non-nested networks, and from 0.24 to 1.0 with R^2^ from 0.26-0.45 in nested networks. With AF, the slopes of the quadratic regressions range from -0.05 to 0.04 with all R^2^ at 0.00 in non-nested networks, and from 0.01-0.07 with R^2^ from 0.00-0.02 in nested networks.

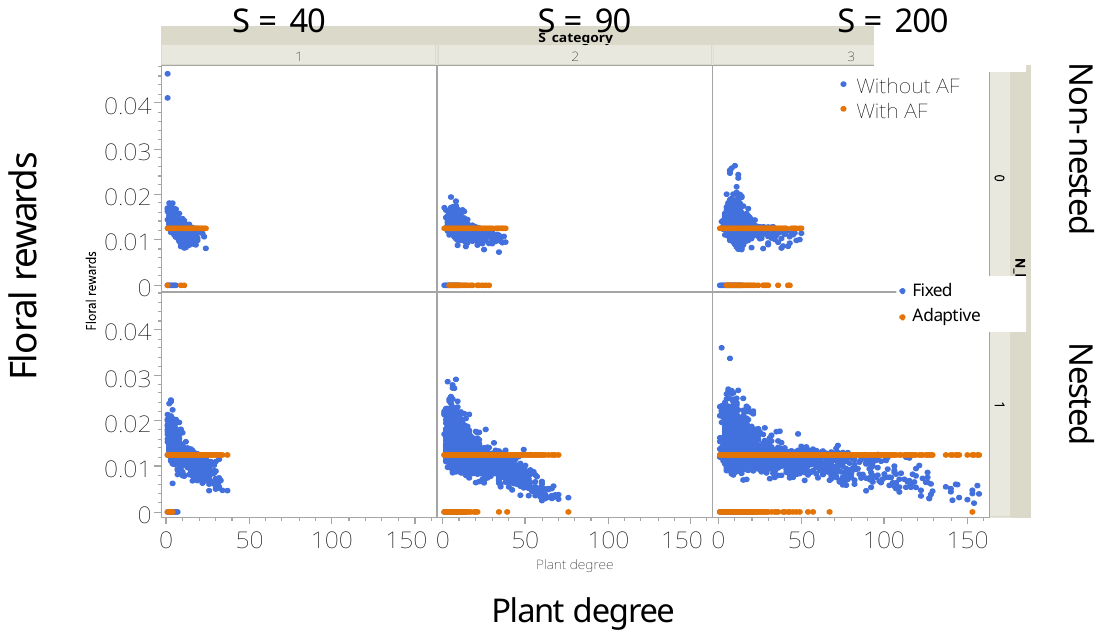

**Fig. S2.** **Final floral rewards with respect of plant species’ degree.** Each dot represent data for each plant species in non-nested (top) and nested (bottom) panels at richness levels (S) of 40, 90, and 200 species; with foxed (blue) and adaptive (orange) foragers. With fixed foragers, generalist plant species contain much lower floral rewards compared to specialist species. With adaptive foragers, floral rewards of all persisting plant species converge to the same value of 0.0125, which is the minimum rewards level pollinators need to persist, or R*.

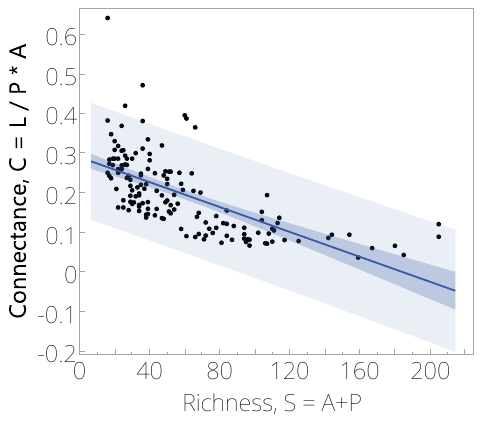

**Fig. S3.** Connectance (C) vs species richness (S) of 155 empirical plant-pollinator networks obtained from <https://www.web-of-life.es/> that contain between 15 and 240 species. Dots represent each of those 155 networks, while the blue solid line, and blue and light blue shaded areas represent the regression between C and S, and its confidence and predicted intervals, respectively. The predicted interval (light blue shade) is used as the boundaries for generating the 1200 networks used in the main text, which are presented in Figure 1A.

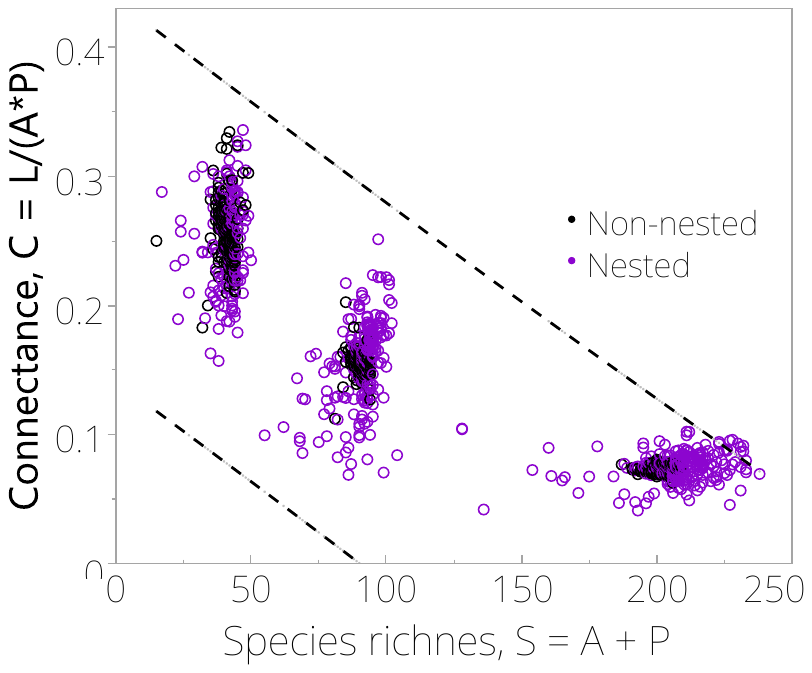

**Fig. S4.** Connectance (C) vs species richness (S) of the 1200 algorithm-generated networks, which are within the 95% prediction intervals of 155 empirical networks with richness between 15 and 240 species (Fig A1). Black and purple circles indicate non-nested and nested network.
