## Supplementary material for "Equalizing Effect of Pollinator Adaptive Foraging on Plant Network Coexistence": All mathematical analyses

### Supplementary Results

#### 1. Niche overlap

From Eqn. (5) in the main text, the per-capita growth of plant species $i \in P$ is given by:

$$\frac{1}{p_i}\frac{dp_i}{dt} = g_i\left(1 - \sum_{l \neq i \in P_j} u_l p_l - w_i p_i\right)\left(\sum_{j \in A} \frac{e_i \alpha_{ij}^2 \tau_j^2 a_j p_i}{\sum_{q \in P} \alpha_{qj} \tau_{qj} p_q}\right) - \mu_i^{(P)}. \tag{S.1}$$

MacArthur (1970) proposed that two species coexist when their intraspecific competition exceeds interspecific competition. The competitive effect of plant species $l$ on $i$ is:

$$\frac{\partial}{\partial p_l}\left(\frac{1dp_i}{p_i dt}\right) = -g_i u_l \sum_j e_i \sigma_{ij} \frac{V_{ij}}{p_i} - \gamma_i \sum_{j \in A_i \cap A_l} \frac{e_i \alpha_{lj} \alpha_{ij}^2 \tau_j^3 a_j p_i}{(\sum_{q \in P} \alpha_{qj} \tau_j p_q)^2}, \tag{S.2}$$

$$\frac{\partial}{\partial p_l}\left(\frac{1dp_i}{p_i dt}\right) = -\left(g_i u_l S_i + \frac{\gamma_i}{p_l} \sum_{j \in A_i \cap A_l} e_i \alpha_{ij} \tau_j a_j \sigma_{ij} \sigma_{lj}\right), \tag{S.3}$$

$$\frac{\partial}{\partial p_l}\left(\frac{1dp_i}{p_i dt}\right) = -\left(g_i u_l S_i + \frac{\gamma_i}{p_l} \sum_{j \in A_i \cap A_l} s_{ij} \Omega_j(i, l)\right). \tag{S.4}$$

$S_i$ and $s_{ij}$ represent $i$'s total seed production and per-capita seed production from pollinator $j$'s visits, respectively. $\Omega_j(i, l)$ is $j$'s visits overlap between plants $i$ and $l$.

The competitive effect of plant species $i$ on itself is:

$$\frac{\partial}{\partial p_i}\left(\frac{1}{p_i}\frac{dp_i}{dt}\right) = -w_i g_i S_i + \gamma_i \sum_{j \in A_i} \frac{e_i \alpha_{ij}^2 \tau_j^2 a_j (\sum_{q \in P - \{i\}} \alpha_{qj} \tau_j p_q)}{(\sum_{q \in P} \alpha_{qj} a_j \tau_j p_q)^2}, \tag{S.5}$$

$$\frac{\partial}{\partial p_i}\left(\frac{1}{p_i}\frac{dp_i}{dt}\right) = \left(-w_i g_i S_i + \frac{\gamma_i}{p_i} \sum_{j \in A_i} s_{ij} \sigma_{ij} (1 - \sigma_{ij})\right), \tag{S.6}$$

$$\frac{\partial}{\partial p_i}\left(\frac{1}{p_i}\frac{dp_i}{dt}\right) = \left(-w_i g_i S_i + \frac{\gamma_i}{p_i} \sum_{j \in A_i} s_{ij} \Omega_j(i, P - \{i\})\right). \tag{S.7}$$

$\Omega(i, P - \{i\})$ is the overlap between $j$'s visits to plant species $i$ and the other plant species in the network. Note that both inter-specific (Eq. S.4) and intra-specific (Eq S.7) competition contain competition for seed recruitment and the effect of pollination via seed production.

Following MacArthur (1970) and Kuang and Chesson (2008), the niche overlap between any pair $(i, l)$ of focal species is obtained as:

$$\rho(i,k) = \sqrt{\frac{\frac{\partial}{\partial p_l}\left(\frac{1}{p_i}\frac{dp_i}{dt}\right) \times \frac{\partial}{\partial p_i}\left(\frac{1}{p_l}\frac{dp_l}{dt}\right)}{\frac{\partial}{\partial p_i}\left(\frac{1}{p_i}\frac{dp_i}{dt}\right) \times \frac{\partial}{\partial p_l}\left(\frac{1}{p_l}\frac{dp_l}{dt}\right)}}. \tag{S.8}$$

Therefore,

$$\rho(i,l) = \sqrt{\frac{(u_l g_i S_i)(u_i g_l S_l) + \frac{\gamma_i}{p_l}(u_i g_l S_l)\sum_j s_{ij}\sigma_{ij}\sigma_{lj} + \frac{\gamma_l}{p_i}(u_l g_i S_i)\sum_j s_{lj}\sigma_{ij}\sigma_{lj} + \frac{\gamma_i\gamma_l}{p_i p_l}\sum_j s_{ij}\sigma_{ij}F_{lj}\sum_j s_{lj}\sigma_{ij}\sigma_{lj}}{(w_i g_i S_i)(w_l g_l S_l) - \frac{\gamma_i}{p_i}(w_l g_l S_l)\sum_j s_{ij}\sigma_{ij}(1-\sigma_{ij}) - \frac{\gamma_l}{p_k}(w_i g_i S_i)\sum_j s_{lj}\sigma_{lj}(1-\sigma_{lj}) + \frac{\gamma_i\gamma_l}{p_i p_l}\sum_j s_{ij}\sigma_{ij}(1-\sigma_{ij})\sum_j s_{lj}\sigma_{lj}(1-\sigma_{lj})}} \tag{S.9}$$

#### 2. Plant fitness

From Eq. (S.1) we have:

$$\frac{1}{p_i}\frac{dp_i}{dt} = g_i\left(1 - \sum_{l \neq i \in P_j} u_l p_l - w_i p_i\right)\sum_{j \in A}\frac{e_i \alpha_{ij}^2 \tau_j^2 a_j p_i}{\alpha_{ij}\tau_j p_i + \sum_{q \neq i \in P}\alpha_{qj}\tau_j p_q} - \mu_i^{(P)} \tag{S.10}$$

Following Kuang and Chesson (2008), we define plant species $i$'s fitness ($k_i$) as its per-capita growth when the competitive effects from itself and other plant species is zero. Thus, we get

$$\Rightarrow \frac{1}{p_i}\frac{dp_i}{dt} = g_i\left(1 - w_i p_i\right)\sum_{j \in A} e_i \alpha_{ij}\tau_j a_j - \mu_i^{(P)}, \tag{S.11}$$

Eliminating the intra-specific effect of the focal species, we obtain per-capita growth of species ($i$) or demographic potential ($k_i$) from Eqn. S.11 as:

$$k_i = g_i \sum_{j \in A} e_i \alpha_{ij}\tau_j a_j - \mu_i^{(P)}, \tag{S.12}$$

$$k_i = g_i e_i v_i - \mu_i^{(P)}. \tag{S.13}$$

where $v_i$ is the total visits plant species $i$ receives per-capita, or total visits per-plant $i$.

#### 3. Plant density at equilibrium

The plant equation is given by:

$$\frac{dp_i}{dt} = g_i \left(1 - \sum_{l \neq i \in P_j} u_l p_l - w_i p_i\right) \sum_{j \in A} e_i \sigma_{ij} V_{ij} - \mu_i^{(P)} p_i. \tag{S.14}$$

$U_i = 1 - \sum_{l \neq i} u_l p_l$ represents $i$'s recruitment potential reduced by inter-specific competition. Plant density at equilibrium ($p_i^*$) can be obtained by solving:

$$0 = p_i \left( g_i (U_i - w_i p_i) \sum_{j \in A} e_i \sigma_{ij} \alpha_{ij} a_j \tau_j - \mu_i^{(P)} \right). \tag{S.15}$$

with two possible solutions: $p_i^* = 0$ ($i$ goes extinct) or solving for the parenthesis equal to zero:

$$0 = g_i \left(U_i - w_i p_i\right) \frac{\sum_{j \in A} e_i \sigma_{ij} \tau_j \alpha_{ij} a_j \sum_{j \in A} \tau_j \alpha_{ij} a_j}{\sum_{j \in A} \tau_j \alpha_{ij} a_j} - \mu_i^{(P)} \tag{S.16}$$

$$0 = g_i \left(U_i - w_i p_i\right) e_i \bar{\sigma}_i v_i - \mu_i^{(P)} \tag{S.17}$$

where $\bar{\sigma}_i = \frac{\sum_{j \in A} \sigma_{ij} \tau_j \alpha_{ij} a_j}{\sum_{j \in A} \tau_j \alpha_{ij} a_j}$ is the per-capita weighted average quality of visit for plant species $i$, and $v_i$ is again total per-plant visits. Solving for $p_i$ in Eq. (S.17) gives us the plant density at equilibrium:

$$p_i^* = \frac{U_i}{w_i} - \frac{\mu_i^{(P)}}{g_i w_i e_i \bar{\sigma}_i v_i} \tag{S.18}$$

The feasibility condition for the equilibrium is $p_i^* \geq 0$, which is obtained when $Q_i = \bar{\sigma}_i v_i$ is higher or equal to:

$$Q_i^c = (\bar{\sigma}_i v_i)^c = \frac{\mu_i^{(P)}}{g_i e_i U_i}. \tag{S.19}$$

Plant species $i$ persists when $Q_i$ is higher than this critical value, while it goes extinct when it is equal or lower than this critical value.
